# *Photorhabdus Luminescens* Toxin complex (TCC) a recombinant injection nano-machine – delivery of recombinant *Yersinia enterocolitica* YopT

**DOI:** 10.1101/698027

**Authors:** Peter Ng’ang’a, Julia K. Ebner, Matthias Plessner, Klaus Aktories, Gudula Schmidt

## Abstract

Engineering delivery systems for proteins and peptides into mammalian cells is an ongoing challenge for cell biological studies as well as for therapeutic approaches. *Photorhabdus luminescens* toxin complex (PTC) is a heterotrimeric protein complex able to deliver diverse protein toxins into mammalian cells. We engineered the syringe like nano-machine for delivery of protein toxins from different species. Additionally, we loaded the highly active copepod luciferase *Metridia longa* M-Luc7 for accurate quantification of injected molecules. We suggest that besides the size also the charge of the cargo defines the efficiency of packing and transport into mammalian cells. Our data show that the *Photorhabdus luminescens* toxin complex constitutes a powerful system to inject recombinant proteins, peptides and potentially other molecules like aptamers into mammalian cells. In contrast to other protein transporters based on pore formation, the cargo is protected from degradation. The system opens new perspectives for cell research and pharmacology.

## INTRODUCTION

*Photorhabdus luminescens* is an entomopathogenic bacterium. It produces a large, heterotrimeric toxin complex (PTC) assembled by three different components: TcA, TcB and TcC which form a syringe-like injection apparatus pre-loaded with a toxin (Sheets & Aktories, 2017, Waterfield, Hares et al., 2005). Homologues of the *Photorhabdus* toxin complex have been identified in other insect and human pathogenic bacteria (Hinchliffe, Hares et al., 2010). Different from the Type-III secretion machineries of *Yersinia spp.* (see below), the *Photorhabdus* megadalton protein complex allows the delivery of its cargo in the absence of the bacteria. Injection of the cargo by the isolated toxin complex requires initial receptor binding on the target cell membrane and depends on structural rearrangements triggered by a shift to higher or to lower pH (Gatsogiannis, Lang et al., 2013, Meusch, Gatsogiannis et al., 2014). The toxin complex is functional on insect, as well as on mammalian cells (Hares, Hinchliffe et al., 2008, Lang, Schmidt et al., 2010). The acidic pH within early endosomes is sufficient to allow toxin injection through the endosomal membrane into the cytosol. The molecular rearrangement of the Tc structure has been analysed in detail by cryo-electron microscopy: TcA is a pentamer forming an alpha helical channel surrounded by a shell-like structure (Gatsogiannis et al., 2013). Channel and shell are connected by a proline rich relaxed peptide, which contracts to a partially helical fold upon cell contact and acidification thereby pushing the channel through the membrane and releasing the pre-loaded toxin into the cytosol (Gatsogiannis, Merino et al., 2016). TcB and TcC together form a cocoon-like structure (BC), which interacts with the TcA pentamer (Meusch et al., 2014, Roderer, Hofnagel et al., 2019). Binding of BC to the pentamer opens a gate, forming a joint channel for loading the toxic enzyme encoded in the hypervariable region (hvr) of the C component (Gatsogiannis et al., 2016). TcC provides the hvr and an aspartyl autoprotease necessary to cleave off the enzyme for its release (Meusch et al., 2014). The modular composition of Tc allows loading and injection of various Photorhabdus toxins into mammalian cells presenting it as a possible vehicle for transport and injection of diverse enzymes and peptides of foreign origin. Here, we engineered the toxin complex to allow delivery of *Yersinia enterocolitica* YopT, a typical Type–III secreted effector protein, into HeLa cells.

*Yersinia enterocolitica* is a food-borne pathogen causing acute and chronic gastro-enteric infections in humans. The bacterium injects several effector proteins into mammalian cells by a syringe-like type-III secretion system presuming the direct contact of the bacteria with the eukaryotic cell (Boyd, Grosdent et al., 2000). The effectors, including the protease YopT, (Yersinia outer protein T) are encoded on a virulence plasmid (pYV) (Iriarte & Cornelis, 1998). YopT is a cysteine protease specifically cleaving the isoprenylated C-terminus of Rho GTPases (Shao, Merritt et al., 2002). Rho proteins are posttranslationally modified at their C-terminal CaaX-box by isoprenylation of the cysteine (C), truncation of -aaX (a-aliphatic, X-any amino acid) and methylation (Zhang & Casey, 1996). This modification is required for localization at cellular membranes and for transport by GDI (Garcia-Mata, Boulter et al., 2011). By removing the isoprenylated cysteine, YopT leads to release of Rho proteins from cellular membranes and liberation from GDI (Aepfelbacher, Trasak et al., 2003, Shao, Vacratsis et al., 2003). Rho proteins are crucial regulators of important signalling pathways ranging from actin-dependent migration and immune cell function to survival (Castellano, Chavrier et al., 2001, Nobes & Hall, 1994). More specifically, YopT blocks phagocytic cup formation, chemotaxis and proliferation (Aepfelbacher et al., 2003, Iriarte & Cornelis, 1998, Trulzsch, Sporleder et al., 2004). Functional injection of several YopT-based proteins allowed us to define the requirements for foreign proteins transported by the Photorhabdus toxin complex. For accurate quantification, we additionally cloned the highly active secreted luciferase of the marine copepod *Metridia longa* as reporter protein into TcBC, because the enzyme meets all pre-conditions for delivery identified in our analysis.

## RESULTS

### Generation of the recombinant toxin chimera BC3-C3bot

It was shown previously that in the toxin complex the enzymatic component to be injected (hypervariable region, hvr) is cleaved within the BC3 cocoon (Meusch et al., 2014). The two toxins of the natural *Photorhabdus luminescens* effector cocktail so far analysed are ADP-ribosyltransferases (TcC3, TcC5). As proof of concept we asked whether we could exchange the hypervariable region of TcC3 towards C3bot (ADP-ribosyltransferase of *Clostridium botulinum*) for functional transport of the *Clostridium* ADP-ribosyltransferase into mammalian cells (Aktories & Koch, 1997). Therefore, C3bot was cloned C-terminally to the auto-proteolytic cleavage site of truncated BC3 (functional cocoon-forming B-C fusion protein of *Photorhabdus luminescens* containing TcB2 and TcC3 (Meusch et al., 2014)) and the chimeric complex was expressed in *Escherichia coli* (Fig. 1A). As expected, the recombinant toxin chimera was auto-proteolytically cleaved (Fig. 1B) into the 25 kDa C3bot and an about 246 kDa fragment (BC-N-term). C3bot ADP-ribosylates the small GTPase RhoA. To study correct folding and activity of the fusion toxin, we performed *in vitro* ADP-ribosylation assays with recombinant RhoA and radiolabelled NAD as second substrate in a time-dependent manner. As shown in Fig. 1C, the fusion toxin was able to modify RhoA. However, compared to C3bot alone (Fig. 1C bottom), activity of the fusion toxin (Fig. 1C top) was much weaker, most likely, because in the chimeric fusion protein C3bot is packed within the cocoon.

**Figure 1:**
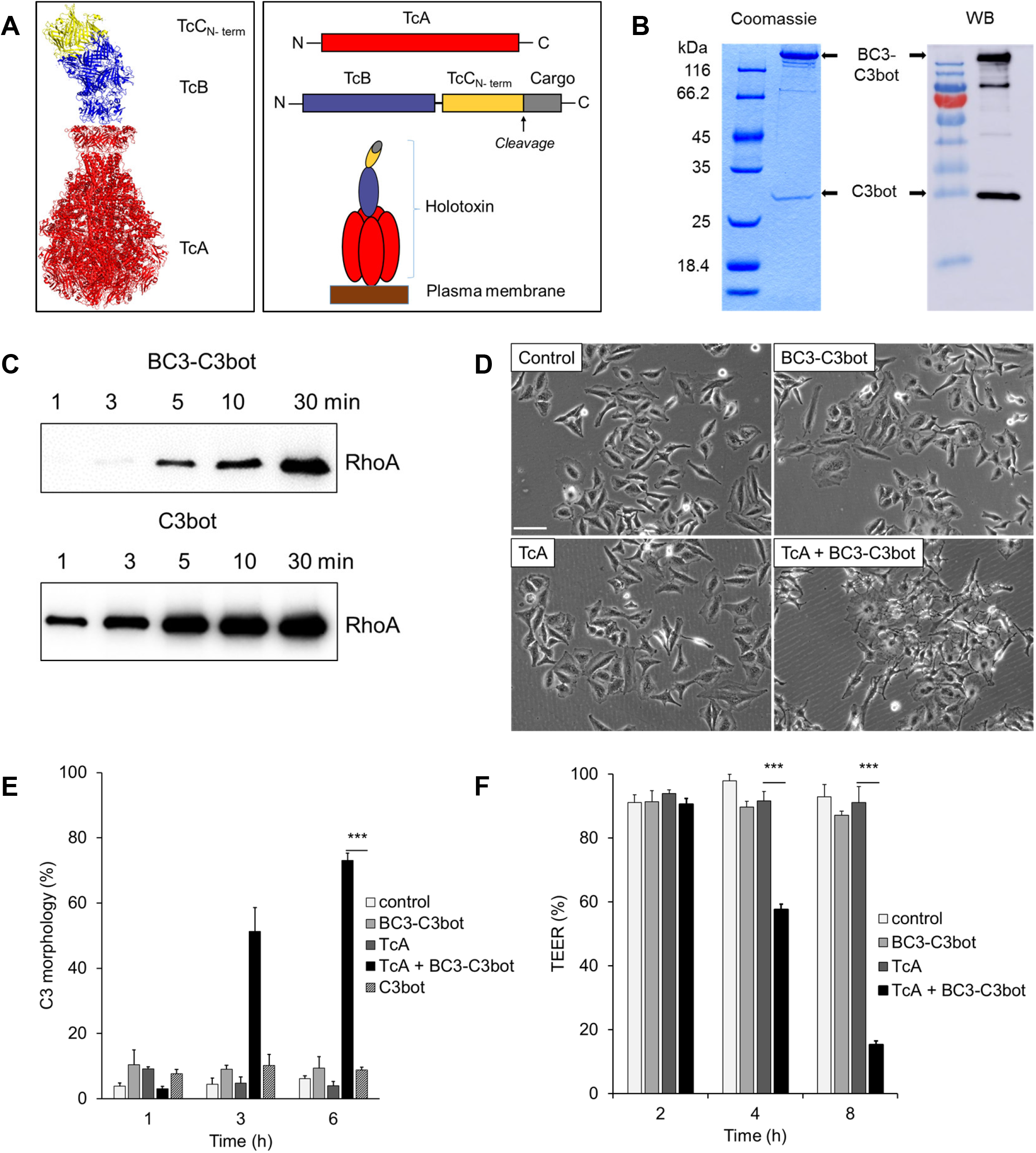
Activity of BC3-C3bot fusion toxin. **A**: Design of the fusion toxin. The TcC3 C-terminal hvr was replaced with a selected cargo following the cleavage site (Adapted from Meusch et al., 2014). **B**: The purified BC3-C3bot fusion protein was separated on an SDS-PAGE gel (Coomassie). For the Western blot (WB), a C3bot-specific antibody was used (Aktories et al., 1989). **C**: Autoradiogram of i*n vitro* ADP-ribosylation of RhoA by 3 nM of BC3-C3bot and 1.25 nM C3bot(WT). **D**: Intoxication of HeLa cells with 10 nM of TcA + BC3-C3bot each for 6 h at 37 °C. **E**: Quantification of intoxication of HeLa cells treated as in D with 100 nM of C3bot(WT). **F**: TEER assay of CaCo-2 cells treated with 1.94 nM of TcA and 5.1 nM of BC3-C3bot for 8 h. Unpaired, two-tailed T-test (P < 0.001) (±SEM). Scale bar: 100 µm. N = 3.

To get insight into the question whether the cocoon shields the toxin from its substrate and/or second substrate, we incubated *Photorhabdus* BC3 at elevated temperatures (4°C, 40°C, 95°C) to destroy the cocoon, cooled the samples down to 21°C again and performed *in vitro* ADP-ribosylation of actin. Interestingly, pre-incubation of BC3 at 95°C led to the highest activity (Fig. S1A) whereas C3hvr alone (Fig. S1B) showed no temperature dependence and was stable up to 95°C. The data indicate that the cocoon shields the catalytic C3 toxin from its substrate. Following artificial opening upon temperature elevation, the heat-stable C3hvr was most likely released.

The most crucial question of the engineered protein injection machinery was its effect on living cells. To analyse effective transport of C3bot by the recombinant toxin complex, HeLa cells were incubated with TcA (*Photorhabdus luminescens* toxin complex subunit TcdA1), BC3-C3bot, TcA plus BC3-C3bot, recombinant C3bot alone, or were left untreated, respectively. As shown in Fig. 1D, only cells treated with both components of the toxin complex rounded up, indicating RhoA inactivation. This proves functional delivery of C3bot by the protein complex into HeLa cells (Fig. 1D, quantification shown in Fig.1E). Analogue to the HeLa experiment, a second cell line was studied. In colon carcinoma cells (CaCo-2 cells) inactivation of RhoA leads to opening of the epithelial barrier reflected by a drop of the trans-epithelial electrical resistance (TEER) (Gerhard, Schmidt et al., 1998). Cells were seeded on filter culture inserts and grown to confluence. Cells were then incubated with each single toxin component or with the full complex, respectively. Following different incubation times, TEER was measured. Consistent with the analyses of HeLa cells, the resistance built by the epithelial barrier exclusively dropped in the presence of the full toxin complex (Fig. 1F). The data show that *Clostridium botulinum* C3 was injected by *Photorhabdus luminescens* toxin complex as foreign cargo into mammalian cells.

### Preconditions for functional delivery

The enclosed tripartite toxin structure carries the active enzymatic component of the toxin sheltered from the environment. Structure analysis revealed a negatively charged inner surface of the cocoon (Gatsogiannis et al., 2016, Gatsogiannis, Merino et al., 2018). This suggests the cargo to be positively charged. To get insight into the features of the cargo, which are required for effective packing and transport, we first compared the properties of naturally appearing substrates of the nano-machine although not all are characterized yet. As shown in Table 1, all putative proteins injected by the *Photorhabdus* toxin complex have a similar molecular weight (between 28 to 35 kDa). Remarkably, their isoelectric point (with one exception) is above 8.5. Our artificial cargo C3bot shows similar characteristics with an even lower molecular weight and a pI of 9.57. The comparison of the open reading frames of possible *Photorhabdus* toxins delivered by PTC and C3bot suggests that the size and the charge of the cargo is key for effective transport. However, so far exclusively ADP-ribosyltransferases and proteins with unknown activities were included in our studies. For further analyses, we chose the *Yersinia enterocolitica* effector protein YopT. The cysteine protease is injected by type-III secretion into mammalian cells to cleave and inactivate Rho GTPases. We generated *Photorhabdus* BC3 fused to YopT proteins of different sizes and isoelectric points (compare Tab.1). Besides full length YopT, we cloned the smaller N-terminal truncations YopTdelta1-30 and YopTdelta1-74 shown to possess full activity and stability when expressed as recombinant protein (Sorg, Hoffmann et al., 2003). The acidic charge of YopTdelta1-74 (pI = 6.2) was varied by C-terminal addition of 3 or 6 lysines, respectively.

**Table 1:** Properties of *Photorhabdus* proteins (in grey) injected by the *Photorhabdus* toxin complex (PTC) and various proteins selected for characterizing the complex’s translocation capabilities

| Gene/Protein name | Activity | MW (kDa) | pI | Reference |
| --- | --- | --- | --- | --- |
| TccC3hvr_W14 | ADP-ribosyltransferase | 32 | 9.68 | Lang *et al.*, 2010 |
| TccC2_W14 | Unknown | 28 | 8.7 |  |
| TccC4_W14 | Unknown | 30 | 6.23 |  |
| TccC5_W14 | ADP-ribosyltransferase | 30 | 8.67 | Lang *et al.*, 2010 |
| TccC1_TT01 | Unknown | 35 | 11.01 |  |
| TccC2_TT01 | Unknown | 28 | 9.4 |  |
| TccC3_TT01 | Unknown | 32 | 9.06 |  |
| TccC4_TT01 | Unknown | 32 | 9.38 |  |
| TccC5_TT01 | Unknown | 30 | 9.08 |  |
| TccC6_TT01 | Unknown | 32 | 8.9 |  |
| TccC7_TT01 | Unknown | 28 | 8.56 |  |
| YopT<sup>Δ1-74</sup> | Cysteine protease | 30 | 6.2 | Shao *et al.*, 2003 |
| YopT<sup>Δ1-74</sup> – lys3 | Cysteine protease | 30 | 8.29 | Shao *et al.*, 2003 |
| YopT<sup>Δ1-74</sup> – lys6 | Cysteine protease | 30 | 8.91 | Shao *et al.*, 2003 |
| YopT<sup>Δ1-30</sup> | Cysteine protease | 30 | 9.04 | Shao *et al.*, 2003 |
| YopT full | Cysteine protease | 36 | 8.53 | Shao *et al.*, 2003 |
| C3bot | ADP-ribosyltransferase | 25 | 9.57 | Aktories *et al.*, 1988 |
| *Metridia* luciferase 7 | Luciferase | 16.5 | 8.66 | Markova *et al.*, 2015 |

### Functional injection of YopT proteins into HeLa cells

We first checked whether the chimeric proteins are catalytically active by analysing the release of RhoA from purified cell membranes. Therefore, membranes were incubated with and without the BC3-YopT proteins, separated into pellet and supernatant by ultracentrifugation and analysed for the presence of RhoA in each fraction. Incubation of the membranes with functional YopT should release cleaved RhoA from the membranes. As shown in Fig. 2A, all constructs showed YopT activity. Heat inactivated enzyme was not able to cleave and release RhoA. Next, we analysed the functional delivery of YopT and YopT truncations by the toxin complex into HeLa cells. Therefore, cells were incubated with the BC3-YopT constructs in the presence or absence of TcA, respectively. Following overnight incubation, cells were fixed and stained for F-actin with rhodamine-phalloidin. As shown in Fig. 2B (top two lanes), artificial injection of YopT led to diminished formation of actin stress fibres, indicating inactivation of RhoA-dependent signalling pathways. Compared to the other constructs, BC3-YopTdelta1-74 had no or only little activity, although it was able to cleave RhoA from purified membranes (Fig. 2A). In contrast, addition of 3 or 6 lysines changing the pI of the cargo led to injection of YopTdelta1-74. In contrast, incubation of cells with TcA only (control) had no effect. As we have shown before, the YopT phenotype could not be reverted by the bacterial toxin Cytotoxic Necrotizing Factor 1 (CNF1) (Sorg, Goehring et al., 2001). CNF1 leads to constitutive activation of Rho proteins by deamidation and strong stress fibre formation. However, mis-localization of the activated Rho protein following YopT injection led to dominant destruction of stress fibres (Sorg et al., 2001). As expected, YopT-induced destruction of stress fibres was even more visible when cells were co-treated with CNF1 (Fig. 2B, bottom two lanes). Again, incubation with BC3-YopTdelta1-74 in the presence of TcA as well as incubation of cells with TcA only had no effect. In contrast to the stress fibres, polymerised cortical actin was still present following YopT delivery. The data indicate that the YopT proteins were injected into HeLa cells and that besides the size of the protein the charge would be essential for functional delivery.

**Figure 2:**
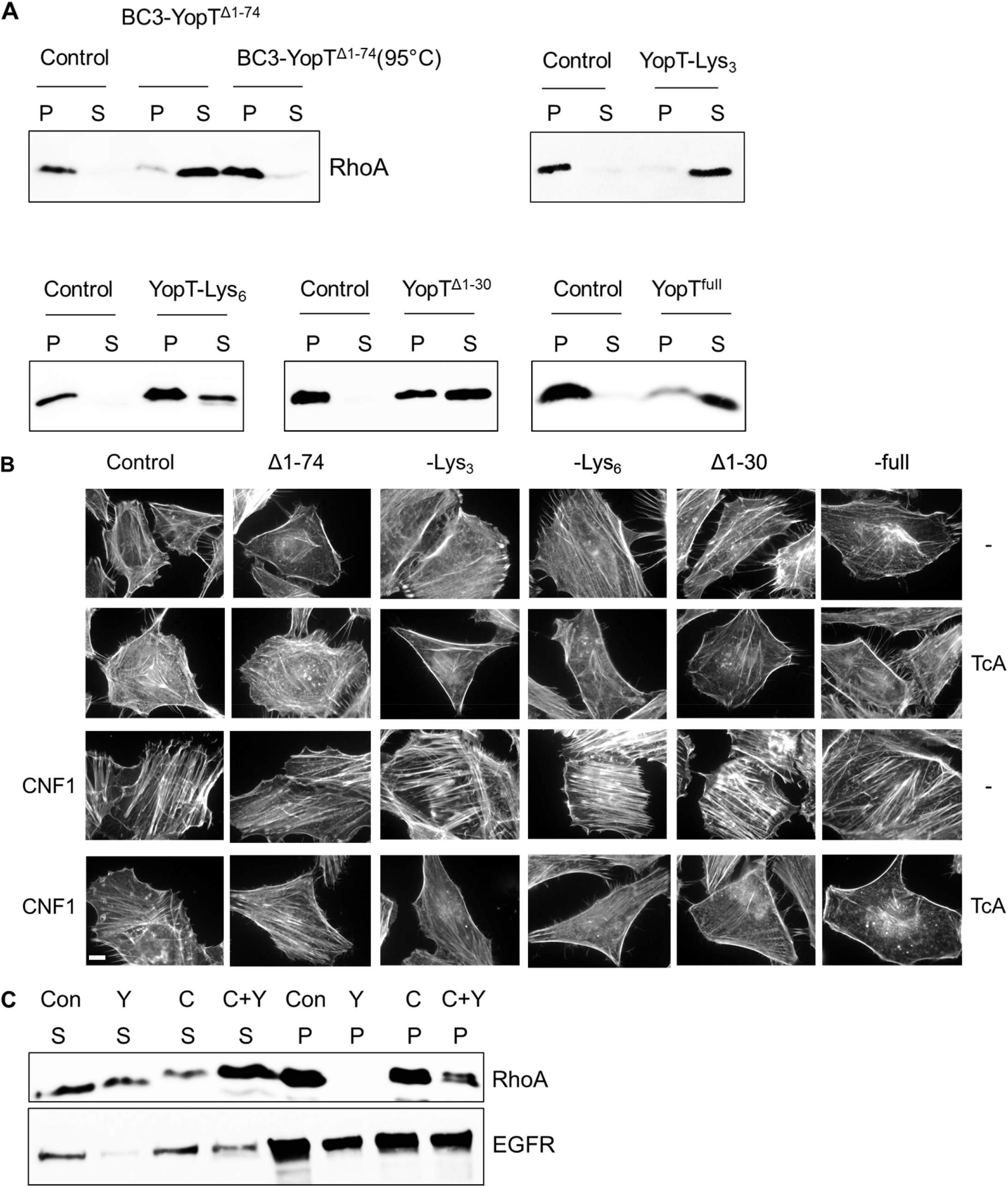
Activity of BC3-YopT fusion toxins. **A**: *In vitro* membrane release activity of RhoA from purified HeLa cell membranes by BC3-YopT chimeras. P = Pellet (membrane fraction), S = post-membrane supernatant. **B**: *In vivo* activity of BC3-YopT. HeLa cells were first intoxicated for 4 h with TcA + BC3-YopTs (20 nM), then 1 h with CNF1 (4 nM) before being fixed and actin stained. **C**: Biochemical analysis of *in vivo* membrane release of RhoA by BC3-YopT^full^ in HeLa cells intoxicated as in B above. P = Pellet (membrane fraction), S = supernatant (cytosolic fraction). Scale bar: 100 µm. N = 3.

For further proof of YopT activity inside the cells, we studied cleavage and mis-localization of RhoA by separation of membranes and cytosol of cells incubated with the respective TC complex proteins (Fig. 2C). These results mirrored the previous ones, with a complete release of RhoA from membranes of YopT-treated cells and a reduction of RhoA release in cells treated with YopT plus CNF1, further confirming successful delivery of YopT into the cells.

### Time resolved analysis of YopT action

As shown before, inactivation of Rho GTPases leads to the destruction of epithelial barriers (Zihni, 2014). However, the effect of recombinant YopT on tight junctions has not been studied because of missing transport into cells. Using the PTC-mediated injection of YopT, we compared the activity of the YopT constructs by analysing their effect on Caco-2 (human coloncarcinoma) cells in a time-resolved manner. Therefore, confluent CaCo-2 cells (TEER = 100%) were treated with increasing concentrations from 1 nM to 30 nM of each toxin chimera, respectively. Transepithelial resistance was measured continuously for up to 20 h. PTC3 was used as positive control and expectedly decreased the epithelial barrier within a few hours. As shown in Fig. 3A, in the presence of TcA, BC3-YopTdelta1-74 did not change the trans-epithelial resistance of the monolayer, which is consistent with the HeLa cell experiments, indicating that YopTdelta1-74 was not actively injected. In contrast, together with TcA all other BC3-YopT constructs induced opening of epithelial junctions with comparable activity (Fig. 3B–E) Fig. 3F shows a direct comparison of all YopT chimeras with a constant concentration of 10 nM. The effective concentration (EC50) of all constructs was similar (15-20nM) with BC3-YopTd30 being the most effective protein (compare Tab. 2, Fig. S2). The data indicate that YopT was injected by the *Photorhabdus* injection machinery into CaCo2 cells. Moreover, YopTwas sufficient to destroy the epithelial barrier by mis-localization of Rho GTPases. *Photorhabdus* toxin complex-based YopT chimeras therefore, may now be used for biochemical analysis of recombinant YopT, because the effector was injected most likely into all cells of a culture. However, the YopT data do not allow for quantification of the injected cargo molecules.

**Figure 3:**
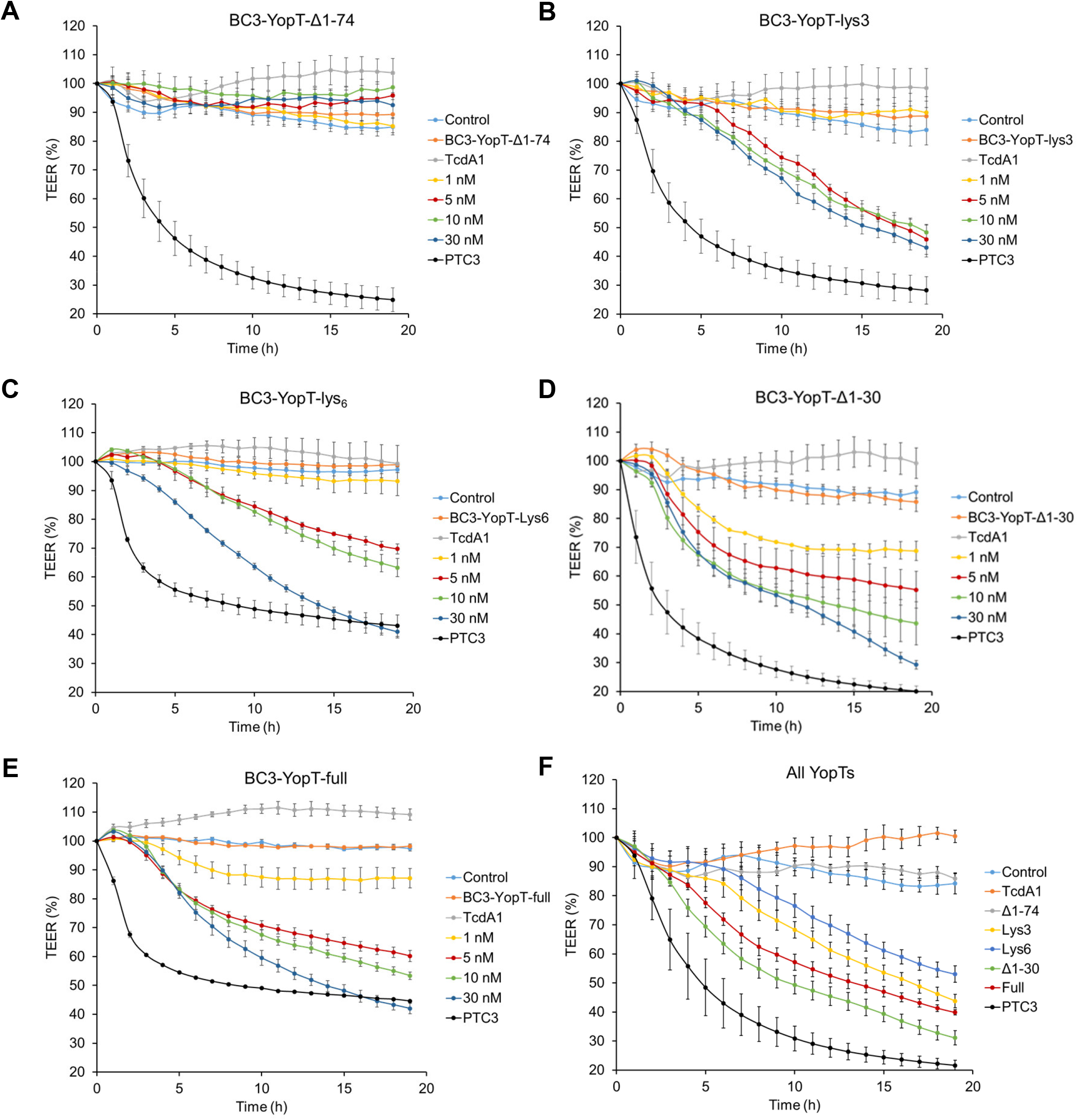
*In vivo* activity of BC3-YopT fusion toxins. Confluent monolayers of CaCo-2 cells were intoxicated with increasing concentrations of TcA + BC3-YopT fusion toxins and TEER was measured. The graphs show TEER as a percentage of the starting value. **A**: BC3-YopT^Δ1-74^. **B**: BC3-YopT^Δ1-74^-lys3. **C**: BC3-YopT^Δ1-74^-lys6. **D**: BC3-YopT^Δ1-30^. **E**: BC3-YopT^full^. **F**: A comparison of all toxins (30 nM each). Recombinant photorhabdus toxin complex 3 (PTC3) was used as a positive control. N = 3.

**Table 2:** Comparison of the EC50 of various BC3-YopT fusion proteins after a 20-hour intoxication of CaCo-2 cells in a transepithelial electrical resistance (TEER) assay

| Toxin | EC50 |
| --- | --- |
| YopT<sup>Δ1-74</sup>-lys3 | 17.43 |
| YopT<sup>Δ1-74</sup>-lys6 | 19.7 |
| YopT<sup>Δ1-30</sup> | 8.27 |
| YopT<sup>full</sup> | 14.52 |

### Quantitative analysis of injected luciferase

Only few molecules of injected toxins are required for complete modification of substrate and the subsequent morphological changes of the cells. Therefore, the highly active toxins are not feasible for exact quantification of the cargo injected into each cell. Based on the composition of the nano-machine, each toxin complex injects a single enzyme into the cell. However, how many PTCs bind to each single cell and functionally release the load remains an open question. To get insights into the number of molecules injected, we made use of a secreted luciferase produced by the marine copepod *Metridia longa* (M-Luc7) which was cloned 15 years ago (Markova, 2004). This enzyme encompasses all requirements needed for functional packing and injection. M-Luc7 (cloned without the signal peptide) has a small molecular weight (152 aa, 16.5 kDa) and a basic character (pI = 8.66). It shares no sequence or structural homology to Renilla or Firefly luciferases (for review see (Markova, Larionova et al., 2019)) although, like Renilla luciferase, it uses commercially available cell penetrating coelenterazine as substrate for producing blue light.

Following binding to mammalian cells, the *Photorhabdus* toxin complex is taken up by endocytosis. Upon intoxication, a higher amount of luciferase is detected when cells have been incubated with the full toxin compared to the cocoon alone, indicating that the luciferase is taken up into the cells (Fig. 4A). However, to distinguish between endocytosis and injection, we incubated the cells at 4°C with only the loaded cocoon (BC3-Mluc7) or with the full toxin complex. Injection was then induced by a pH change to acidic pH (pH 5) or the cells incubated at neutral pH 7.5 as indicated. Cells were washed and treated with trypsin/EDTA to degrade extracellular protein and simultaneously to detach the cells from the dish. Cells were harvested, counted, lysed and lysates transferred into a white plate for detecting luciferase activity. Similarly, at both pH 5 and 7.5, a higher amount of luciferase was detected when cells were incubated with the full toxin compared to the cocoon alone, indicating that the luciferase was delivered into the cells (Fig. 4B). In acidic pH however, much more luciferase was measured, suggesting favourable injection of the enzyme at this pH (Fig. 4B, compare Fig. 4D and E). Interestingly, comparing endocytic and injection-based delivery of YopT, the latter revealed much higher activity, especially at pH 5 (Fig. 4A and B). In line with this, injection at the plasma membrane closely mimics the native system where the effectors are injected via the type-III secretion system.

**Figure 4:**
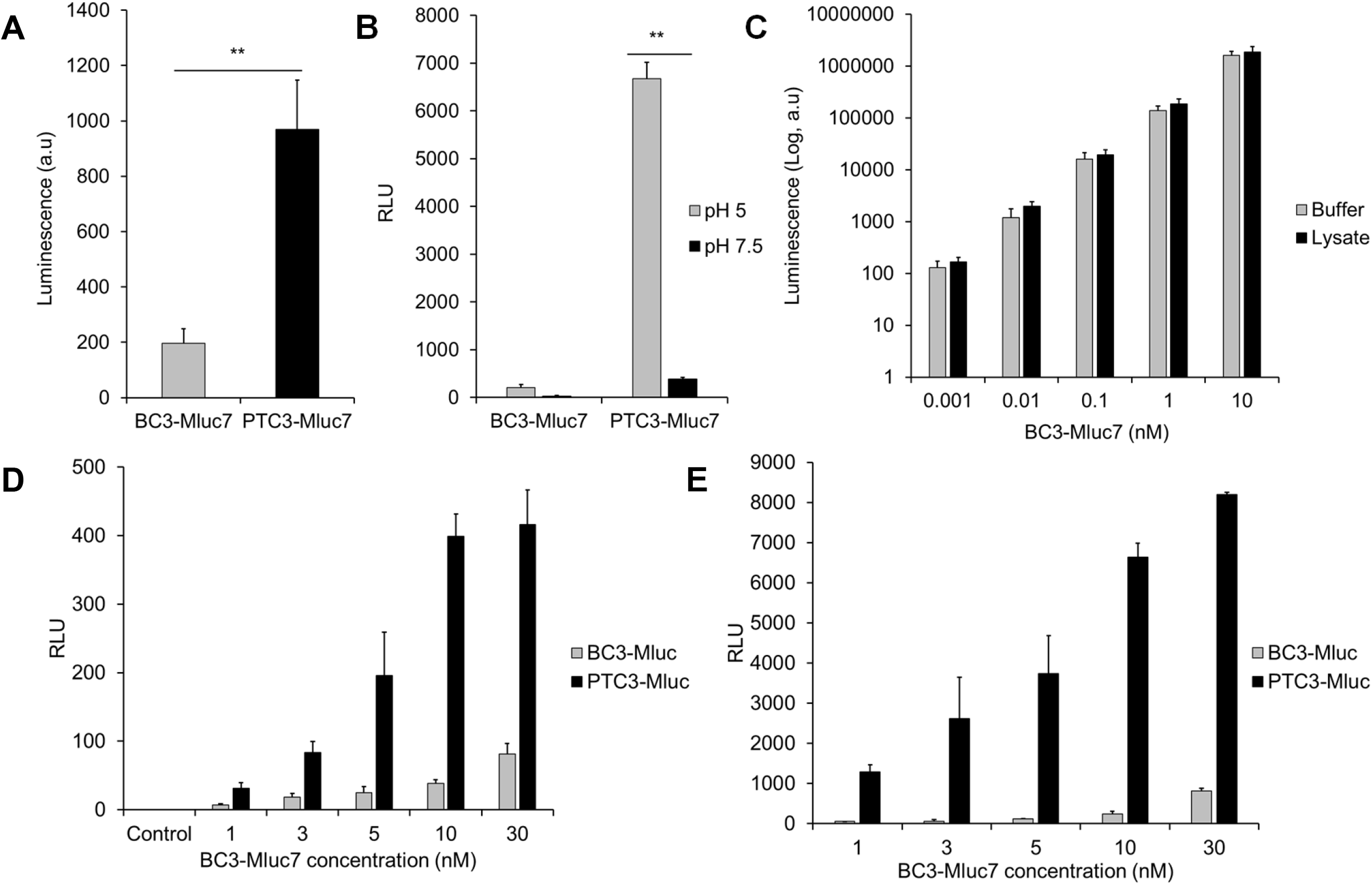
Luciferase activity of BC3-Mluc7 fusion toxin. A: *In vivo* activity of Mluc7 after treatment of HeLa cells with 10 nM of TcA + BC3-Mluc7 for 1 h at 37 °C. B: pH-dependent delivery of MLuc7 across the cell membrane of HeLa cells. RLU = Relative luminescence units (Luminescence/total protein concentration). C: *In vitro* activity of BC3-Mluc7 in passive lysis buffer (Buffer) or HeLa cell lysate. D: pH-dependent delivery of MLuc7 across the cell membrane at pH 7.5. E: pH-dependent delivery at pH 5. Unpaired, two-tailed T-test (P < 0.01) (±SEM). N = 3.

To calculate the injected molecules per cell, a standard curve detecting luminescence produced by increasing amounts of purified BC3-M-Luc7 in buffer and cell lysate was prepared (Fig. 4C, Fig. S3). Additionally, standard curves detecting luminescence produced by injecting M-Luc7 into cells from the cell membrane at pH 7.5 (Fig. 4D) and pH 5 (Fig. 4E) were prepared. From these curves, we estimated about 3.4×10^8^ molecules M-Luc7 produced one light unit in our system (lysate). This corresponds to 30.5 light units in 10^6^ cells produced by 10^10^ molecules M-Luc7, giving 10^4^ molecules injected into each cell at pH 7.5 (961.5 light units giving 3.1×10^5^ molecules per cell at pH 5) when cells were incubated with 1 nM of the full complex.

To further prove intracellular localization of the injected luciferase, we incubated CaCo2-cells with TcC-M-Luc7 (10 nM), buffer, TcA or BC3-M-Luc7, respectively, removed extracellular protein by treatment with trypsin/EDTA, added cell-permeable coelenterazine and detected luminescence by live-cell microscopy using sensitive GaAsP detectors. As shown in Fig. 5, only cells treated with the full complex show a bright blue luminescence, indicating successful delivery of the luciferase.

**Figure 5:**
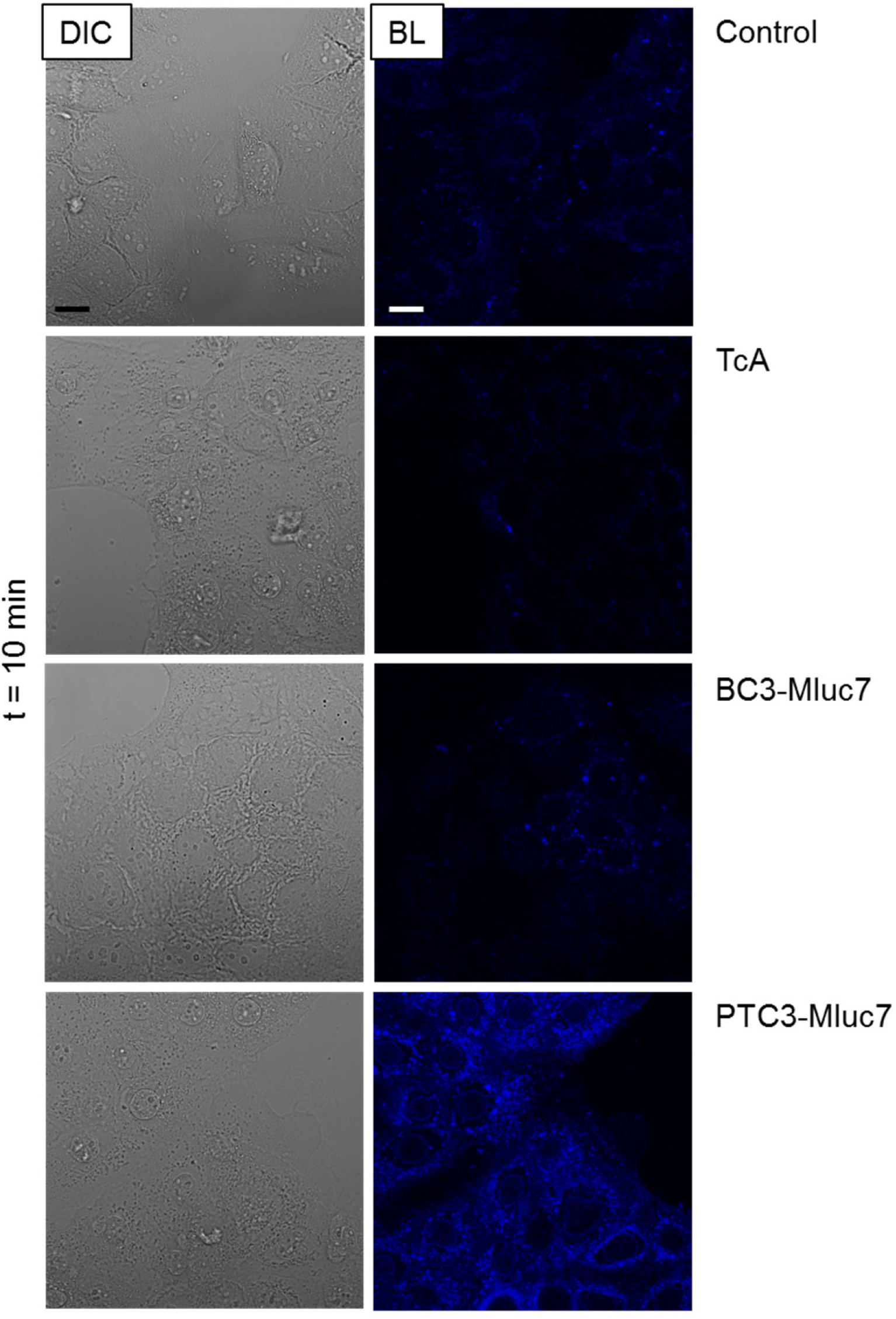
Bioluminescence activity of BC3-Mluc7 fusion toxin. Membrane delivery of toxin (10 nM) into HeLa cells was first done. Then, 2 µg/ml of coelenterazine H was added and the cells visualized for 10 minutes. The controls include coelenterazine only, TcA + coelenterazine, BC3-Mluc7 + coelenterazine. Figure represents one of two independent experiments. DIC = Differential Interference contrast, BL = Bioluminescence. Scale bar: 15 µm.

## DISCUSSION

Transport of proteins through cellular membranes, either for cell biological studies or for therapeutic approaches is a technical challenge for scientists. Besides manual microinjection into single cells, some methods to transport proteins through biological membranes have been developed: Cell penetrating peptides like Antennapedia homeodomain protein and HIV-1 Tat are able to cross cellular membranes and have been used to transport small proteins fused to it into cells (Rizzuti, Nizzardo et al., 2015), although the exact mechanism of membrane passage is not known. Moreover, peptidomimetics were developed to mimic the conformation of peptide sequences. However, mimicking larger protein surfaces is not yet possible (deRonde & Tew, 2015). Pore forming bacterial toxins like *Bacillus anthracis* protective antigen or *Clostridium botulinum* C2 toxin have been exploited to deliver recombinant proteins across cell membranes (Barth, Roebling et al., 2002, Collier & Young, 2003). Bacteria are able to actively inject cargo proteins into mammalian cells by type-III secretion machineries (Ballard, Collier et al., 1996, Cornelis, 2000, Mechaly, McCluskey et al., 2012). Such bacteria have been genetically manipulated as delivery vector for vaccination, which requires the direct contact of living bacteria with mammalian cells (Derouazi, Wang et al., 2010, Epaulard, Toussaint et al., 2006, Nishikawa, Sato et al., 2006). Here, we characterized a standalone injection nanomachine of the entomopathogenic bacterium *Photorhabdus luminescens*. Physiologically, it injects different proteins into insect cells, most of them with unknown function (compare Tab.1). Two of the injected toxins however are already characterized as ADP-ribosyltransferases modifying actin and Rho GTPases, respectively (Lang et al., 2010). We showed that the *Clostridium botulinum* ADP ribosyltransferase C3bot was accepted as foreign cargo. When injected by the nanomachine, the toxin induces typical cell rounding within 6 h by destruction of the actin cytoskeleton. In former studies, C3, which consists of an enzymatic domain exclusively, was delivered by pore forming toxins like *Clostridium botulinum* C2 toxin. Using this kind of delivery system, uptake of C3 led to cell rounding within 2 to 3 h (Barth et al., 2002), suggesting a more effective transport. However, an advantage of the PTC over the pore forming transporters may be the protection from degradation within the closed cage.

To study whether also other enzymes than ADP-ribosyltransferases would be packed and injected, we further fused the protease *Yersinia enterocolitica* YopT to BC. YopT is physiologically injected by Type-III secretion. It localizes to cellular membranes and releases mainly RhoA by cleaving off the C-terminal isoprenylated cysteine (Shao et al., 2003). Moreover, RhoA is released from GDI, leading to proteasomal degradation of the small GTPase and to disruption of stress fibres (Aepfelbacher et al., 2003, Zumbihl, Aepfelbacher et al., 1999). In our study, YopT was released from endosomes. In contrast to manual microinjection or expression of YopT, experiments in which cell rounding occurred within minutes, destruction of stress fibres and opening of barriers was much slower when only few molecules of YopT were injected. This may more closely reflect the physiological situation, including substrate specificity. Although purified YopT does not show specificity for any specific Rho GTPase *in vitro*, spatial localization of the injected protease may be the basis for its specificity in living cells (Sorg et al., 2001). In mammalian cells, RhoA is attached to the plasma membrane, whereas Rac and Cdc42 show endosomal and Golgi localization (Michaelson, Silletti et al., 2001). In *Yersinia-*infected macrophages, RhoA seems to be the preferred substrate of YopT (Aepfelbacher et al., 2003). In our experiments using the *Photorhabdus* toxin complex for injection of YopT, cells do not round up completely. However, stress fibre formation was reduced, even in the presence of the Rho activating toxin CNF1. (In former experiments, YopT was injected manually, cells rounded up within minutes, which could not be reverted by CNF1 (Sorg et al., 2001).

In *Yersinia*, the N-terminal amino acids of YopT are involved in binding of its chaperone SycT (Buttner, Cornelis et al., 2005). Chaperone interaction is required for efficient translocation by the type-III secretion system (Trulzsch et al., 2004). However, the truncated recombinant proteins showed activity and high stability in *in vitro* assays (Sorg et al., 2003). Therefore, YopT truncations were analysed for injection by the *Photorhabdus* toxin complex. The fragment with the highest stability and *in vitro* activity showed no or very little effects on HeLa- and Caco-2 cells. Two reasons could be raised: unfolding of the packed cargo may be necessary for loading or the basic pI un-favours loading into TcA. Therefore, we changed the pI by adding 3 or 6 lysines which brought back activity, indicating that an acidic pI was essential for successful injection.

According to the assembly of the injection apparatus, each nanomachine is capable of injecting a single molecule. We therefore asked how many toxin complexes do bind to each single cell and are able to inject their load. To answer this crucial question, we made use of the marine luciferase *Metridia longa* M-Luc7. The enzyme is released from the copepod to glare its predator. It does not need ATP but converts coelenterazine into light by oxidation. Because in living cells are reducing conditions, we would expect lower activity compared to buffer conditions. Our calculations therefore may be too low. Truly, our data can give only a rough estimate. However, we calculated about 10^4^ molecules of MLuc7 injected into each cell at pH 7.5 and 3.1×10^5^ molecules per cell at pH 5 when cells were treated with the full complex (concentration 1nM). This leads us to suggest that 0.5% of the luciferase entered cells (0.016% at pH 7.4).

With this highly sensitive and quantifiable injected luciferase at hand, we will be able to compare the efficiency of further engineered toxin complexes to get insight into efficient binding and pore formation. By engineering the *Photorhabdus luminescens* toxin complex, we developed a powerful system to inject proteins, peptides and potentially other molecules like aptamers into mammalian cells opening new perspectives for cell biological and therapeutic approaches.

## MATERIALS AND METHODS

### Materials

Cell culture medium DMEM and fetal calf serum were purchased from Biochrom (Berlin, Germany), whereas McCoy’s 5A medium was purchased from PAN Biotech (Aidenbach, Germany). Cell culture materials were obtained from Greiner (Frickenhausen, Germany). Ni-IDA resin was from Macherey-Nagel (Düren, Germany). [^32^P]NAD^+^ and ^86^RB were from PerkinElmer (Cologne, Germany). *Clostridium histolyticum* collagenase type IA was purchased from Sigma (now Merck, Darmstadt, Germany). Bafilomycin A1 was purchased from Enzo life sciences (Lörrach, Germany). Coelenterazine H (Biotium) was purchased from Hölzel Diagnostika (Cologne, Germany).

### Protein expression and purification

Proteins were expressed and purified as previously described (Lang et al., 2017). In short, *E. coli* BL21 (DE3) cells were transformed with *P. luminescens* TccC3hvr and protein expression was induced by the addition of IPTG to a final concentration of 75 µM. For *P. luminescens* TcdA1, TcdB2-TccC3 or TcdB2-TccC3 fusion proteins, *E. coli* BL21-CodonPlus cells were transformed and protein expression was induced with 25 µM IPTG. After 24 h, all *P. luminescens* protein-expressing cells were harvested and resuspended in lysis buffer (300 mM NaCl, 20 mM Tris-HCl, pH 8.0, 1 mM DTT, 500 µM EDTA, and 10% glycerol) supplemented with DNase (5 µg/ml), lysozyme (1 mg/ml) and 1 mM PMSF. After sonication, cell lysate was incubated with Ni-IDA resin and loaded onto empty PD-10 columns. The His6-tagged proteins were eluted with 500 mM NaCl, 20 mM Tris-HCl, pH 8.0, 0.05% Tween-20, 500 mM imidazole, and 5% glycerol. The protein-containing fractions were pooled and dialyzed against 100 mM NaCl, 50 mM Tris, pH 8.0, 0.05% Tween-20, and 5% glycerol. Purification of fusion toxins was confirmed by Western blotting with specific antibodies (Aktories, Braun et al., 1989).

For cleavage of TcdA1 by collagenase, the protein was first incubated with collagenase (3 µg of TcdA1 with 150 ng (~93 nM) of collagenase) for 1 h at 37°C in TcdA1 storage buffer without glycerol. Then, the cleaved toxin was separated from collagenase by fast protein liquid chromatography (FPLC) in TcdA1 storage buffer without glycerol as previously described (Ost et al., 2018). This involved size-exclusion chromatography with ÄKTApurifier through a Superose 6 Increase column (GE healthcare, Freiburg, Germany).

### Cell culture and cytotoxicity assays

HeLa and CaCo-2 cells were cultivated at 37°C and 5% CO_2_ in DMEM containing 10% heat-inactivated FCS, 2 mM L-glutamine, 0.1 mM non-essential amino acids, 100 units/ml penicillin, and 100 µg/ml streptomycin. HT-29 cells were cultured in McCoy’s 5A medium supplemented with 10% FCS and penicillin/streptomycin as mentioned above. For cytotoxicity experiments, cells were seeded in culture dishes and incubated in medium with 0.5% FCS together with the respective toxins. After the indicated incubation periods, cells were visualized using a Zeiss Axiovert 40CFI microscope (Oberkochen, Germany) with a Jenoptik Progress C10 CCD camera (Jena, Germany). The cytopathic effects caused by the toxins were analyzed in terms of morphological changes and quantified by counting the number of intoxicated cells.

### Transepithelial electrical resistance (TEER) assays

For the TEER assay, the electrical cell-substrate impedance sensing (ECIS) system (Applied biophysics, New York, USA) was used. CaCo-2 cells were seeded on 8-well 8W10E+ ECIS arrays (Ibidi GmbH, Martinsried, Germany) and cultured for 2 days. Assays were performed when TEER values remained constant at ~1000-1600 ohms (100% confluence). First, the arrays were precooled on ice for 15 min. Then, indicated toxins were added and allowed to bind for 1 h. Finally, the arrays were transferred to 37 °C where TEER was measured. The *Photorhabdus* toxin complex 3 (PTC3, 0.7 nM TcA + 1.75 nM BC3) was used as a positive control.

### ADP-ribosylation of actin and RhoA

For *in vitro* ADP-ribosylation of actin, 1.25 nM of TccC3hvr or 3 nM of BC3, was incubated with 1.9 µM βγ-actin, 150 µM NAD^+^, radioactive [^32^P]NAD^+^ (0.5 μCi per sample) at 21°C in buffer containing 5 mM Hepes, pH 7.5, 0.1 mM CaCl_2_, 0.1 mM ATP and 0.5 mM NaN_3_. After the indicated time points, the reaction was stopped by addition of SDS-containing Laemmli buffer. Then, the samples were subjected to SDS-PAGE and radiolabeled actin was detected and visualized by autoradiography. For *in vitro* ADP-ribosylation of RhoA, 60 nM of recombinant wildtype C3bot or BC3-C3bot fusion protein, was incubated with 0.4 µM GST-RhoA, 1 µM NAD^+^, radioactive [^32^P]NAD^+^ (0.5 μCi per sample) and 2 µg BSA at 37°C in buffer containing 25 mM TEA, 2 mM MgCl_2_, 1 mM DTT and 1 mM GDP. After the indicated time points, the reaction was stopped by addition of SDS-containing Laemmli buffer, the samples subjected to SDS-PAGE, and radiolabeled RhoA detected and visualized by autoradiography.

### Membrane-release assays

For *in vitro* membrane release assays, HeLa cells cultured in 10 cm dishes were washed once with PBS and then detached by scrapping in 500 µl of lysis buffer (50 mM Tris-HCl, pH 7.4, 150 mM NaCl, 5 mM MgCl_2_, 1 mM EDTA, 1 mM PMSF and 2.5 mM DTT). The cells were then lysed by sonication and clarified by centrifugation at 1,000 × g for 10 minutes at 4 °C. Afterwards, membranes and cytosol were separated by ultracentrifugation at 32,000 rpm for 1 h at 4 ° C. The subsequent pellet, consisting of the membrane fraction, was retained while the supernatant was discarded. This pellet was then resuspended in lysis buffer and either treated with 0.5 µM of each BC3-YopT fusion protein for 30 min at 37 °C or left untreated and just incubated. The samples were then ultracentrifuged again to separate the pellets (membranes) from the supernantants containing released Rho GTPases (post-membrane supernatant). Finally, samples were mixed with SDS-containing Laemmli buffer, separated by SDS-PAGE, transferred onto PVDF membranes and blotted using a RhoA-specific antibody (#2117, Cell signaling technology, Frankfurt, Germany), followed by an HRP-linked anti-rabbit secondary antibody (#7074, Cell signaling technology, Frankfurt, Germany).

For *in vivo* membrane release assays, HeLa cells were first intoxicated with 20 mM of TcdA1 and BC3-YopT^full^ for 4 h at 37 °C, followed by 1 h with 4 nM of CNF1. Afterwards, the cells were washed, lysed, clarified and the lysates ultracentrifuged as described above to separate membranes (pellet) from cytosol (supernatant). The pellets were then immediately resuspended in Laemmli buffer, while the supernatants were precipitated by addition of trichloroacetic acid (1:9 ratio), incubation for 30 min at 4 °C, followed by centrifugation at 14,000 rpm for 5 min. The pellets from these precipitated supernatants were then resuspended in Laemmli buffer and together with the membrane fraction, run on SDS-PAGE and blotted as above.

### Inhibition of cellular stress fibers

Four sets of Hela cells were cultured on coverslips overnight. Then, the cells were washed with PBS and the medium changed to DMEM containing 0.5% fetal calf serum. Afterwards, two sets of cells were intoxicated for 4 h at 37 °C, with 20 nM of TcdA1 together with all the indicated BC3-YopT fusion proteins. Then, one set of YopT-treated cells and an untreated set were intoxicated with 4 nM of CNF1 for 1 h. After CNF1 intoxication, the cells were washed twice with PBS, fixed with 4% paraformaldehyde for 15 min, washed again with PBS and permeabilized with 0.15% (v/v) Triton X-100 for 10 min. Subsequently, the cells were incubated in the dark with phalloidin–tetramethylrhodamine (TRITC) for 2 h at RT for actin staining. Finally, after washing and treatment with 70% and 100% ethanol, respectively, the cells were dried and embedded with Mowiol supplemented with DABCO (1,4-Diazabicyclo[2.2.2]octane) and DAPI (4′,6-diamidino-2-phenylindole) for nuclear staining. Samples were cured for 24 h and the cells visualized using an Axiophot system (Zeiss, Oberkochen, Germany).

### Bioluminescence assays

For *in vitro* bioluminescence assays, the indicated concentration of BC3-Mluc7 was added to 200 µl of either Passive lysis buffer (Promega, USA), HeLa cell lysate, or indicated buffer. Then, this BC3-Mluc7 mix was aliquoted into a white plate and 30 µl of activity buffer (1M NaCl, 20 mM MgSO_4_, 0.03% gelatin and 100 mM Tris HCl, pH 7.4) (Markova, 2004) were added. Finally, the plate was transferred to an infinite M200 microplate reader (Tecan, Austria), where 30 µl of Stop&Glo solution (Promega, USA), diluted 1:1 with Passive lysis buffer, was dispensed into each well and the resultant luciferase activity detected.

For *in vivo* assays, HeLa cells cultured in 10 cm dishes were detached, counted and 500,000 cells/ml taken in PBS for each treatment. The cells were incubated for 1 h at 37 °C with 10 nM of either the loaded cocoon only (BC3-Mluc7) or with the full toxin complex (PTC3-Mluc7; TcA + BC3-Mluc7). Afterwards, the cells were collected by centrifugation for 5 min at 300 × g, washed, resuspended in 200 µl of passive lysis buffer and lysed by incubating on a rotary shaker for 30 minutes at RT. The protein concentration of the lysate was then determined by Bradford assay, after which it was aliquoted into a white plate, activity buffer added and luciferase activity detected as described above.

For pH-dependent assays, HeLa cells were first incubated with 100 nM of Bafilomycin A1 for 30 min at 37 °C. Then, they were transferred to ice, precooled to 4 °C and then incubated for 1 h with either BC3-Mluc7 only or PTC3-Mluc7 in HBSS buffer (1× HBSS, 20 mM Hepes, pH 7.5) at the indicated concentrations. Then, to induce injection of Mluc7 into the cells by the surface-bound toxin complex, the pH of the surrounding medium was changed either to acidic pH 5 medium (medium with 0.5% FCS and 20 mM MES, pH 5), or neutral pH 7.5 (medium with 0.5% FCS and 20 mM Hepes, pH 7.5) as indicated. These cells in pH medium were incubated for 1 min at 37 °C before the medium was shifted back to pH 7.5. Afterwards, the cells were washed and treated with trypsin/EDTA for 2 min at 37 °C to degrade extracellular protein and simultaneously detach them from the plate. Finally, the cells were harvested, lysed in passive lysis buffer and transferred into a white plate for detecting luciferase activity.

For bioluminescence microscopy, CaCo-2 cells were treated as in the pH-dependent assays described above without Bafilomycin A1. In contrast, these cells were not detached by the trypsin/EDTA treatment and were thus able to be visualized by a Zeiss LSM 800 microscope (Oberkochen, Germany). After intoxication with 10 nM of BC3-Mluc7 only or PTC3-Mluc7, 2 µg/ml of coelenterazine H diluted in PBS was added and image acquisition was started. To allow simultaneous bioluminescence and DIC imaging, the 405-nm laser was set to 0.005 mW. Bioluminescence images (between 450 and 700 nm) were acquired with a GaAsP detector set to maximum gain. Images were taken every 30 s for a total duration of 30 min. As controls, substrate only, TcdA1 only and full toxins with YopT^Δ1-74^ and YopT^full^ were used.

### Statistical analysis

To determine the relative luminescence units (RLU), luminescence values were divided by the total protein concentration of the respective lysates. To determine the number of luciferase molecules delivered per cell by the *Photorhabdus* toxin complex, light units emitted from a single cell were calculated. For this, data from *in vitro* bioluminescence assays was used to perform a linear regression with D’Agostino & Pearson omnibus normality test using GraphPad Prism version 5.00 for Windows (GraphPad Software, San Diego California USA). This generated the concentration of BC3-Mluc7 that produces one light unit (LU) and the requisite number of protein molecules. Next, from the pH-dependent assays, luminescence values obtained from 1 nM of PTC3-Mluc7 were used to calculate the respective number of luciferase molecules delivered in total. A separate quantification of the number of cells per well narrowed this value down to light units emitted from a single cell, and from that, the number of luciferase molecules delivered per cell.

## Acknowledgement

We thank Alexander Lang for fruitful discussion and Jürgen Dumbach for excellent technical support.

This work was supported by the Deutsche Forschungsgemeinschaft SFB 850 (project C2 to GS). Financial support of P.N. by the DAAD is gratefully acknowledged.

---

**Figure S1:**
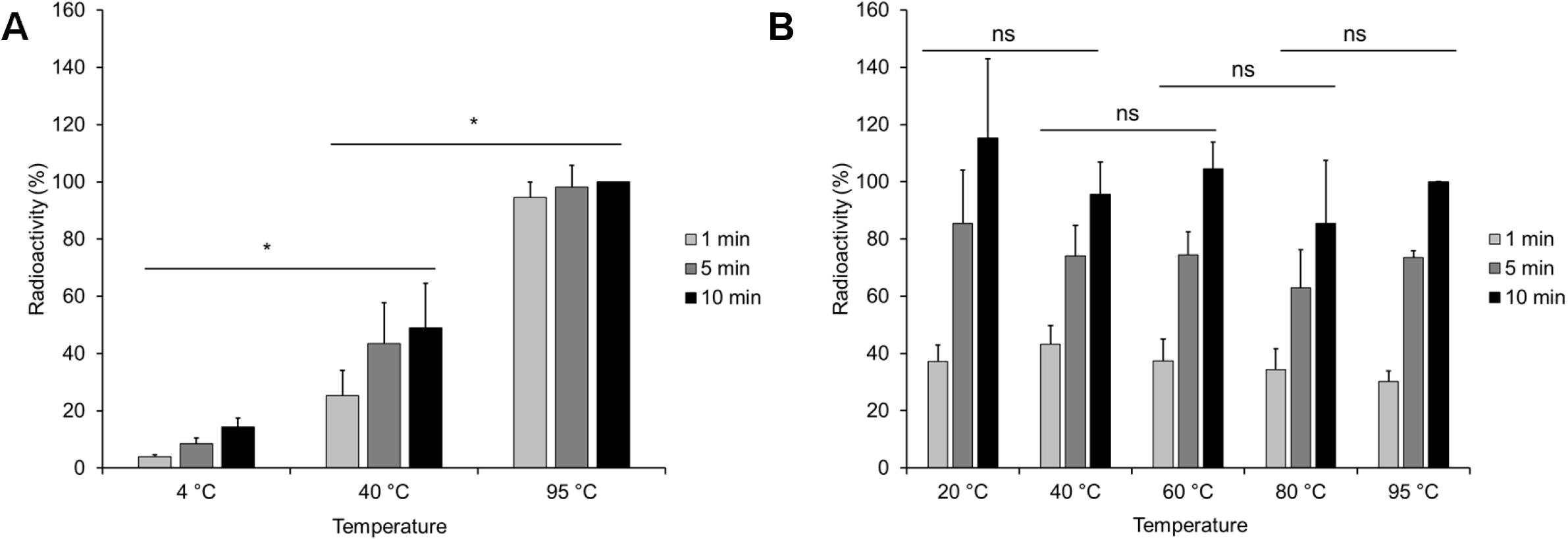
Effect of temperature on the BC3 cocoon. Samples were incubated at the indicated temperatures for 1, 5 and 10 minutes, cooled and the assay conducted at 21 °C. **A**: BC3 (WT) **B**: TccC3hvr. Unpaired, two-tailed T-test (P < 0.05) (±SEM). N = 3.

**Figure S2:**
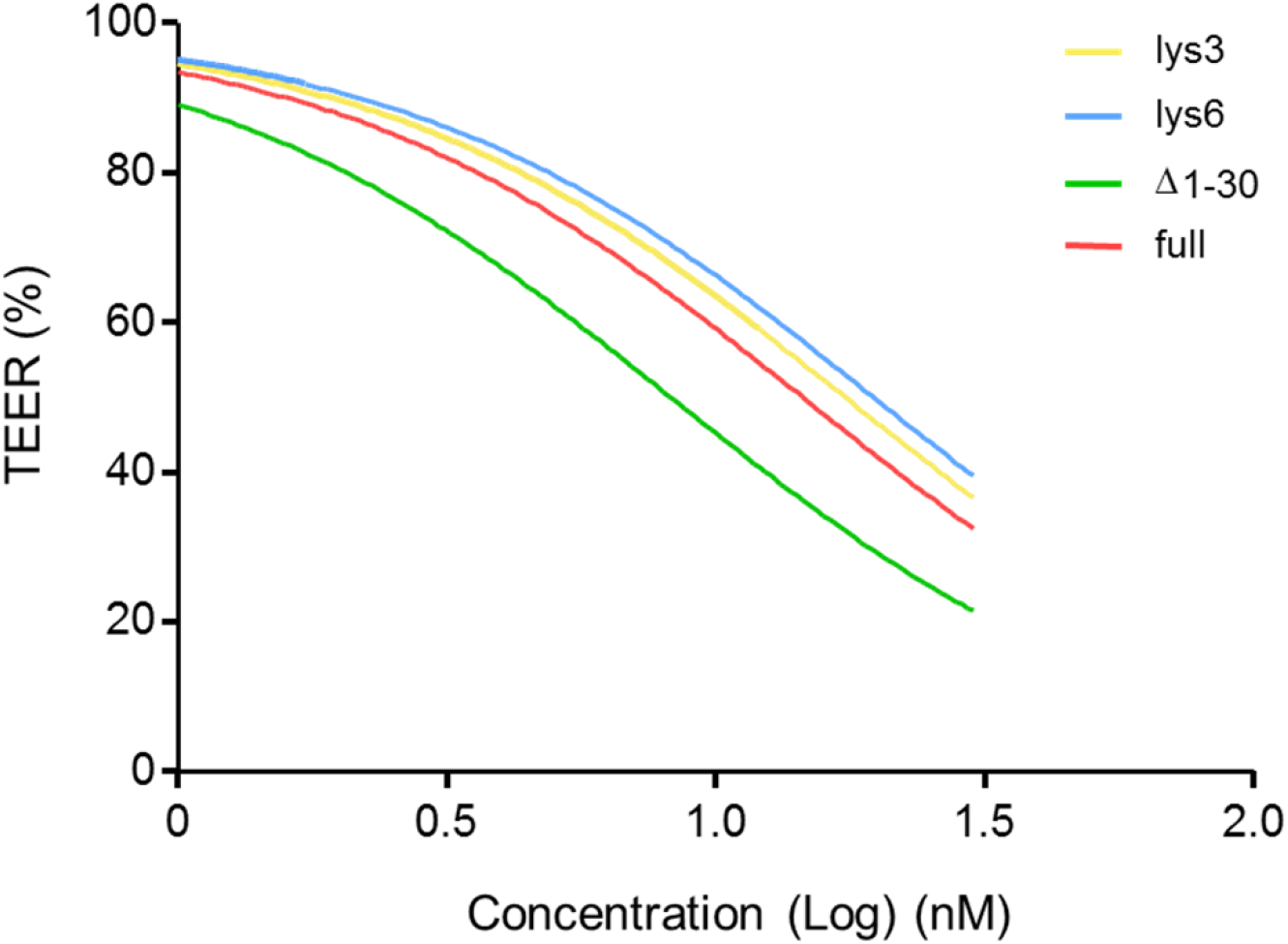
EC50 calculation of BC3-YopT fusion toxins after intoxication of CaCo-2 cells with increasing concentrations of TcA + BC3-YopTs for 20 h. BC3-YopT^Δ1-74^-lys3 (yellow) = 17.43, BC3-YopT^Δ1-74^-lys6 (blue) = 19.7, BC3-YopT^Δ1-30^ (green) = 8.27, BC3-YopT^full^ (red) = 14.52. N = 3.

**Figure S3:**
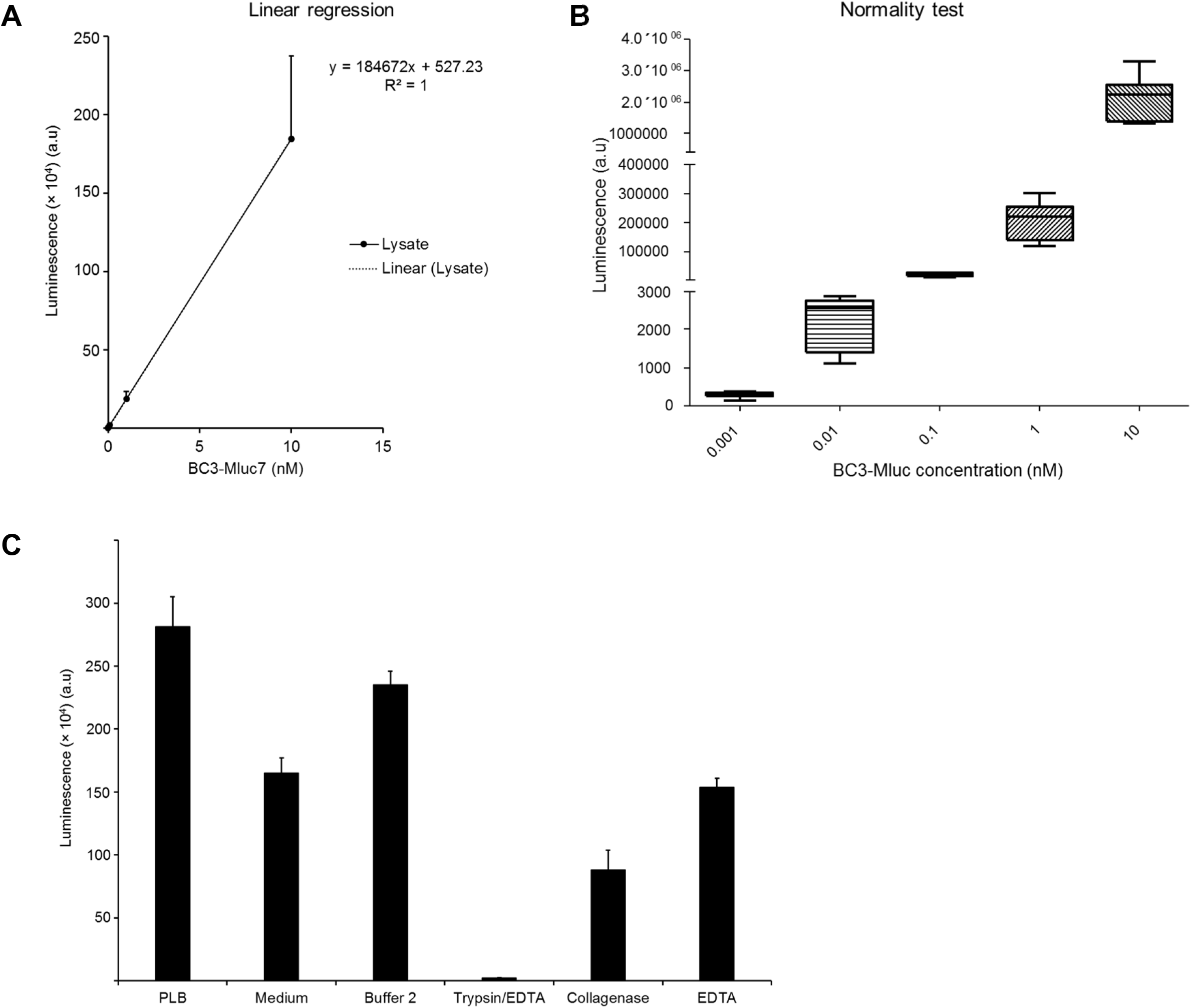
BC3-Mluc7 activity and quantification of the efficiency of PTC3-Mluc7. A: Linear regression calculation of BC3-Mluc7 luciferase activity in HeLa cell lysate. B: Normality test of data used in A. C: *In vitro* activity of BC3-Mluc7 in different buffers and enzyme treatments. ±SEM. N = 3.

